# CellMAGE: cell-type deconvolution for multi-parent population analysis of gene expression

**DOI:** 10.64898/2026.09.08.750196

**Authors:** Robyn L. Ball, Alyssa Klein, Ashley A. Auth, Daniel A. Skelly, Hao He, Vivek M. Philip, Leona Gagnon, Elissa J. Chesler

**Author notes:** Contributed equally to this work (Robyn L. Ball, Alyssa Klein).

## Abstract

Single-cell RNA-sequencing remains prohibitively expensive for multiparental population (MPP) studies. Existing deconvolution methods treat bulk RNA-seq as genetically anonymous mixtures, but in MPPs, the proportional contribution of each parental strain to each progeny’s transcriptome is already known. CellMAGE (Cell-type deconvolution for Multi-parent Analysis of Gene Expression) weights parental cell-type profiles by each progeny’s known genetic composition, requiring no model training and no minimum sample size. Validated in 16 Diversity Outbred mice across 12 prefrontal cortex cell types and 23,116 genes, predicted and measured gene expression were statistically equivalent (±0.05) in all cell types (pooled Spearman ρ = 0.923, 95% CI: 0.910, 0.934). CIBERSORTx required 96 additional samples to resolve at most 12.4% of genes and only 3 cell types; CellMAGE outperformed it even within this restricted comparison (per-cell-type median ρ: 0.908-0.974 vs. 0.353-0.641). CellMAGE is applicable to any MPP with parental single-cell data, including diploid crop MAGIC populations.

**Article summary:** Identifying which cell types manifest genetic effects on complex traits requires cell-type-specific gene expression data, but single-cell sequencing is prohibitively expensive for population genetic sample sizes. Ball et al. present CellMAGE, a computational method that delivers single-cell fidelity from bulk sequencing in multiparental populations, at bulk sequencing cost. Because each individual’s genotypes are known, CellMAGE weights parental cell-type profiles by genetic composition directly, requiring no model fitting. Validated in Diversity Outbred mice across 12 brain cell types, CellMAGE predictions were highly accurate, outperformed an existing method, and apply to any multiparental population with available parental single-cell data, including crop species.

## Introduction

Single-cell RNA-sequencing (scRNA-seq) has transformed our ability to characterize cellular diversity, revealing molecular mechanisms of disease, the functional organization of neural circuits, and the interactions among cell types within complex tissues at unprecedented resolution. The power of single-cell approaches is further amplified in genetically diverse populations, where cell-type-specific expression can be linked to genetic variation to identify the cellular mechanisms underlying complex traits (Yazar et al., 2022; Nathan et al., 2022).

Advanced multiparental populations, including Diversity Outbred (J:DO) and Collaborative Cross (CC) mice, multiparental rat populations, and crop populations produced through the Multiparent Advanced Generation Inter-Cross (MAGIC) design, provide the allelic diversity and combinatorial genetic variation needed to map genetic influences on gene expression and complex traits at high resolution (Svenson et al., 2012; Complex Trait Consortium, 2004; Hansen & Spuhler, 1984). Applied to multiparental populations, single-cell resolution would transform bulk expression quantitative trait loci (eQTL) discoveries into cell-type-resolved genetic maps, revealing the precise cellular contexts through which genetic variation shapes complex phenotypes. Yet large-scale scRNA-seq and snRNA-seq in multiparental populations remain out of reach for most studies, as per-sample costs substantially exceed those of bulk RNA-seq and the large cohort sizes required for genetic mapping amplify this disparity.

Deconvolution methods that predict cell-type-specific gene expression from bulk RNA-seq data offer a scalable alternative, providing cellular resolution without the expense of single-cell profiling. Despite their promise, widespread adoption of these methods remains challenging due to their statistical and computational complexity, minimum sample size requirements, and the expertise required for implementation. More fundamentally, existing deconvolution methods were not designed for multiparental population architectures (Newman et al., 2019; Wang et al., 2019; Jew et al., 2020). These methods rely on reference-based model fitting that treats the reference as a generic single-cell atlas, without accounting for the defining feature of multiparental populations: the fact that each individual’s genome, and therefore transcriptome, is a quantifiable mixture of a small number of known parental genomes. This structure means that the cellular composition of any progeny bulk sample is not unknown, as general deconvolution methods assume, but mathematically predictable from parental genetics. Existing methods cannot exploit this information and instead treat multiparental progeny as they would any heterogeneous bulk sample, discarding the precise genetic knowledge that defines these populations. To our knowledge, no deconvolution method has been developed specifically for multiparental populations that explicitly leverages this known parental genetic structure.

Here we present CellMAGE (Cell-type deconvolution for Multi-parent Analysis of Gene Expression), a simple, accurate, and accessible deconvolution method designed specifically for multiparental populations. CellMAGE exploits the additive genetic architecture of multiparental populations, in which each progeny’s transcriptome reflects a quantifiable, strain-specific mixture of parental contributions, to reconstruct cell-type-specific gene expression from bulk RNA-seq data. The method takes as input two data sources: cell-type-specific gene expression profiles from parental strains, derived from founder snRNA-seq or scRNA-seq, and allele-specific bulk expression estimates for each progeny generated by EMASE (Expectation-Maximization for Allele Specific Expression; Raghupathy et al., 2018; https://github.com/churchill-lab/emase). EMASE is an expectation-maximization algorithm that exploits the hierarchical structure of parental alleles to partition bulk gene expression by founder strain of origin, returning strain-specific expression estimates and total expression for each individual and gene. CellMAGE uses these strain-specific weights to compute a biologically grounded, genetically precise weighted sum of parental cell-type profiles, yielding predicted cell-type-specific transcriptomes per individual. Sex-specific predictions are supported by incorporating same-sex parental expression profiles, enabling accurate prediction in studies with sex as a biological variable. Because CellMAGE derives predictions directly from the known genetic composition of each progeny rather than fitting a statistical model, it requires no model training, imposes no minimum sample size, and performs equally well for a single sample as for a large cohort.

We validate CellMAGE in the prefrontal cortex of 16 Diversity Outbred mice across 12 cell types in a full factorial sex-by-treatment design, demonstrating statistical equivalence of predicted and measured gene expression (±0.05) across all 12 cell types with a pooled Spearman correlation of 0.923. CellMAGE substantially outperforms CIBERSORTx in gene expression fidelity and cell-type coverage, resolving all 20,000+ expressed genes across all 12 cell types compared to at most 3,084 genes in 3 of 12 cell types for CIBERSORTx. CellMAGE is applicable to any multiparental population with available parental single-cell data, including CC mice, F1 crosses, multiparental rat populations, *Drosophila* (Synthetic Population Resource; King et al., 2012), *C. elegans* multiparental recombinant inbred lines (Noble et al., 2017), yeast multiparental populations, and crop diploid MAGIC populations such as in maize, sorghum, rice, and barley (Kumar et al., 2023; Cavenaugh, et al., 2008), as demonstrated in a companion maize (*Zea mays*) example (Miculan et al., 2021; Supplementary Methods). To maximize accessibility, CellMAGE is freely available as an interactive R Shiny application for use without programming expertise and as an R package for large-scale computational pipelines, with full documentation and example datasets at https://thejacksonlaboratory.shinyapps.io/CellMAGE/ and https://github.com/TheJacksonLaboratory/chesler-lab/tree/main/CellMAGE/.

## Methods

### Animals and tissue collection

Parental samples were collected from eight inbred mouse strains (founder strains): A/J (J:000646), C57BL/6J (J:000664), 129S1/SvImJ (J:002448), NOD/ShiLtJ (J:001976), NZO/HlLtJ (J:002105), CAST/EiJ (J:000928), PWK/PhJ (J:003715), and WSB/EiJ (J:001145). Progeny data were collected from Diversity Outbred (J:DO) mice, genetically unique individuals derived from the founder strains (Table 1). One male and one female per founder strain (16 total) and 16 J:DO mice (8 female, 8 male) were tested on consecutive days of behavioral phenotyping (open field, light-dark box, hole board, and novel place preference) at 8 to 10 weeks of age. After behavioral phenotyping, J:DO mice were randomly assigned to cocaine or saline treatment groups in a full factorial experimental design (4 J:DO per sex per treatment); all founder samples came from a repeated saline treatment group used as a control in a cocaine sensitization procedure (Schoenrock et al., 2020). Prefrontal cortex (PFC) tissue was collected from the left-brain hemisphere of all mice following rapid decapitation 24 hours after the final injection of cocaine or saline. Samples were cryo-preserved in liquid nitrogen and stored at −80 degrees Celsius until sequencing (Table 1). Standardized protocols were followed for all drug exposure and sample collection procedures (Chesler et al., 2021; https://phenome.jax.org/projects/CSNA03/protocol). All procedures were approved by The Jackson Laboratory Institutional Animal Care and Use Committee (IACUC protocol 10007) prior to the start of the study.

**Table 1.** Experimental design sample summary.

| Strain / population | Sex |  | Treatment |  | Sequencing |  |
| --- | --- | --- | --- | --- | --- | --- |
|  | Female | Male | Saline | Cocaine | snRNA-seq | Bulk RNA-seq |
| A/J | 1 | 1 | 2 | — | 2 | — |
| C57BL/6J | 1 | 1 | 2 | — | 2 | — |
| 129S1/SvImJ | 1 | 1 | 2 | — | 2 | — |
| NOD/ShiLtJ | 1 | 1 | 2 | — | 2 | — |
| NZO/HILtJ | 1 | 1 | 2 | — | 2 | — |
| CAST/EiJ | 1 | 1 | 2 | — | 2 | — |
| PWK/PhJ | 1 | 1 | 2 | — | 2 | — |
| WSB/EiJ | 1 | 1 | 2 | — | 2 | — |
| **Subtotal**<br>**Founder Strains** | **8** | **8** | **16** | **—** | **16** | **—** |
| Diversity Outbred | 8 | — | 4 | 4 | 8 | 8 |
| Diversity Outbred | — | 8 | 4 | 4 | 8 | 8 |
| Diversity Outbred | 48 | — | 24 | 24 | — | 48 |
| Diversity Outbred | — | 48 | 24 | 24 | — | 48 |
| **Subtotal**<br>**Diversity Outbred** | **56** | **56** | **56** | **56** | **16**<br>**validation set** | **112** |
| **Total** | **64** | **64** | **72** | **56** | **32** | **112** |

Green rows = Founder Strains, serving as cell-type expression input | Purple rows = Diversity Outbred mice; serving as bulk expression input and cell-type expression validation. Numbers indicate N per group.; — = not applicable. All samples: prefrontal cortex, left hemisphere. Founders: saline treatment only. J:DO: full factorial design (sex × treatment).

### RNA-sequencing

Individual samples were collected from whole tissue samples from the left half of the PFC of 16 founders and 16 J:DO mice. To construct the validation set, each J:DO tissue sample was split such that one portion was sequenced using snRNA-seq and the remainder was sequenced in bulk (bulk RNA-seq), in a paired design (Table 1). An additional 96 J:DO bulk RNA-seq samples, balanced for sex and treatment, were included to benchmark CellMAGE against CIBERSORTx, which requires a larger sample cohort to estimate cell-type fractions and gene expression; however, cell-type-specific gene expression was only recoverable for a subset of cell types (Table 1; SRA accessions PRJNA1517767, PRJNA1516495, PRJNA1518853).

#### Single-nucleus RNA-sequencing

Frozen PFC tissue was lysed and then homogenized using a Miltenyi gentleMACS dissociator, following the nuclei isolation protocol (Perry et al., 2022). A 100 µl aliquot of the lysate was immediately removed and placed on ice then stored in a freezer until bulk RNA-sequencing (see bulk RNA sequencing, below). The remaining lysate was immediately processed to isolate nuclei, which were cleaned using a sucrose gradient to remove debris prior to sequencing.

Nuclei viability was assessed on a LUNA FX7 automated cell counter (Logos Biosystems), and up to 30,000 nuclei from each suspension were loaded onto one lane of a 10x Genomic Chip M. Single cell capture, barcoding and library preparation were performed using the 10x Genomics Chromium X platform and version 3.1 Next GEM Single Cell 3’ HT chemistry according to the manufacturer’s protocol (CG000416 Rev D). cDNA and libraries were checked for quality by Tapestation 4200 (Agilent) and Qubit Fluorometer (ThermoFisher), quantified by KAPA qPCR, and sequenced on an Illumina NovaSeq 6000 S4 v1.5 200 cycle flow cell lane, with a 28-10-10-90 asymmetric read configuration, targeting 15,000 barcoded nuclei with an average sequencing depth of 50,000 reads per nucleus.

##### Single-nucleus RNA-sequencing analysis

Separate preprocessing analyses were conducted on snRNA-seq samples from two distinct cohorts: 16 founders (two per sex per strain) and 16 J:DO (4 per sex per treatment). Cell-type-specific gene expression from the founder strains was used as input to CellMAGE. Cell-type-specific gene expression from the 16 J:DOs served as the validation set. Importantly, the validation set was not used at any stage to develop CellMAGE. Each preprocessing analysis followed the procedure described below.

The same preprocessing, quality control (QC), normalization, clustering, and annotation pipeline described below was applied independently to the 16 saline-treated founders and 16 J:DO samples (saline- or cocaine-treated, treatment covariate).

Individual libraries were processed using 10x Genomics Cell Ranger (Zheng et al., 2017) then for each cohort, Cell Ranger output was aggregated into a single Seurat R object (Hao et al., 2021). Outlier nuclei were detected and filtered within sample using median absolute deviation (MAD) in the Scuttle R package (McCarthy et al., 2017) using sample-specific thresholds as the distributions of the number of detected genes and unique molecular identifiers (UMIs) varied across samples; a shared cutoff would have been either too permissive for high-depth samples or too stringent for low-depth samples. Genes and UMIs were excluded if the natural log transformed values were greater than 3 times the MAD from the median. Mitochondrial percentage is expected to be uniformly low across nuclei in snRNA-seq, therefore, a fixed threshold was applied across samples, excluding nuclei with a mitochondrial percentage above 1.5%.

To account for sequencing depth differences within sample, retained nuclei were log-normalized with the Seurat *LogNormalize* method, where UMI counts for each nucleus were divided by the total UMI counts for that nucleus, multiplied by a scale factor of 10,000, and natural log-transformed after adding a pseudo-count of 1, retaining the 2,000 most variable genes across the nuclei. To remove nuisance variation unrelated to cell identity, gene expression residuals from per gene linear regression models were input to principal component analysis (PCA). The first 100 principal components (PCs) were initially retained. An elbow plot of the first 100 PCs revealed that most of the variance (76.8%) was explained by the first 30 PCs. Nuclei were clustered jointly across samples with the first 30 PCs of each sample. A shared nearest neighbor (SNN) graph was constructed across the first 30 PCs, using 20 nearest neighbors per nucleus, and nuclei were clustered using the Louvain algorithm at a resolution of 0.5 (Hao et al., 2021). Clustering visualization was performed using UMAP.

Per cohort, batch effects were corrected with the Harmony R package (Korsunsky et al., 2019). PCA was performed on harmony-based nuclei embeddings then clustered jointly across samples with the first 30 PCs.

Cell types were identified with ScType automated annotation using the scaled gene expression from the Seurat RNA assay and a curated brain marker gene dataset (Ianevski et al., 2022). In addition to the ScType brain cell type marker genes, *Satb2* was added as a glutamatergic neuron marker, as it is predominantly expressed in excitatory neurons of the neocortex (Huang et al., 2013). Cell type scores were computed per nucleus and summed per cluster; the cell type with the highest aggregate score per cluster was assigned as the cluster label. Clusters were flagged as low-confidence and relabeled ‘unknown’ if their top-scoring cell type’s aggregate score fell below a threshold of one-quarter of the total number of cells in that cluster, following the default ScType recommendation (Ianevski et al., 2022). Cells that were not differentiated into a cell type, and were assigned to an ‘unknown’ cluster, were removed from downstream analyses.

Sample and cell type level metrics were computed from the Seurat object metadata. Cells were grouped by sample and cell-type annotation as received by ScType. For each sample/cell type pair we calculated the number of cells (*n_cells*), the mean and median of the number of detected genes per cell (*nFeature_RNA*), total UMI counts per cell (*nCount_RNA*), percent mitochondrial reads (*percent.mt*), and percent ribosomal reads (*percent.ribo*). A gene was considered ‘expressed’ in the sample/cell type if the gene expression was nonzero (count > 0) in at least one cell using the raw count matrix summed across all cells.

##### Cell-type-specific gene expression and cell type fraction

Cell-type-specific (pseudobulk) gene expression matrices were generated for each cell type using Seurat’s AggregateExpression() function. These matrices included columns for Ensembl gene identifier (id), each founder or J:DO sample id, gene expression, as well as meta-data indicating sex and treatment. Pseudobulk gene expression was based on aggregating raw UMI counts across all nuclei annotated to the cell type per sample. Schwann precursor cells are not a component of the central nervous system and due to the low number of observed annotated nuclei, Schwann precursor cells and cells that could not be differentiated (‘unknown’) were excluded from analysis (GEO accession GSE346204).

Cell-type fractions of each J:DO sample were computed from ScType cell annotations. The cell type fraction was calculated as the proportion of the number of cell-type-specific nuclei relative to the total annotated nuclei in the J:DO sample. Nuclei annotated to Schwann precursor cells and unknown cells were excluded prior to computing cell-type fractions.

#### Bulk tissue RNA-sequencing

Total RNA was isolated from the PFC sample of each J:DO (Table 1) using the NucleoMag RNA Kit (Macherey-Nagel) and the KingFisher Flex purification system (ThermoFisher). Tissues were lysed and homogenized in nuclei extraction buffer (Miltenyi) and protector RNase inhibitor (Roche) using a gentleMACS dissociator (Miltenyi Biotec Inc). After an addition of MR1 buffer and TCEP (Macherey-Nagel) samples were mixed by pipetting and RNA isolation was performed according to the manufacturer’s protocol. RNA concentration and quality were assessed using the Nanodrop 8000 spectrophotometer (Thermo Scientific) and the RNA ScreenTape Assay (Agilent Technologies).

RNA-seq libraries were constructed using the KAPA mRNA HyperPrep Kit (Roche Sequencing and Life Science), according to the manufacturer’s protocol. Briefly, the protocol specifies isolation of polyA containing mRNA using oligo-dT magnetic beads, RNA fragmentation, first and second strand cDNA synthesis, ligation of Illumina-specific adapters containing a unique barcode sequence for each library, and PCR amplification. The quality and concentration of the libraries were assessed using the D5000 ScreenTape (Agilent Technologies) and Qubit dsDNA HS Assay (ThermoFisher), respectively, according to the manufacturers’ instructions.

Libraries were sequenced 150 bp paired-end on an Illumina NovaSeq 6000 using the S4 Reagent Kit v1.5.

##### Bulk tissue RNA-sequencing analysis

Prior to alignment, we built the multi-way alignment index of the eight founder strains by introducing strain-specific SNPs and short indels from the Mouse Genome Project release v8 (Keane et al. 2011; https://ftp.ebi.ac.uk/pub/databases/mousegenomes/REL-2112-v8-SNPs_Indels/) to the reference genome GRCm38 Ensembl v78 primary assembly using g2gtools (http://churchill-lab.github.io/g2gtools). Raw paired-end reads from each sample were aligned to the multi-way index with Bowtie v1.3.1 (Langmead et al., 2009).

Because J:DO mice are genetic mosaics of eight founder strains, reads may align to multiple founder genomes, and within a genome, to multiple transcripts of the same gene. J:DO allele-specific expression was estimated from the raw read alignment counts using the Expectation-Maximization algorithm for Allele Specific Expression (EMASE, v0.10.16; Munger et al, 2014), an algorithm originally developed to analyze J:DO gene expression by accounting for the founder strain alleles. EMASE leverages the hierarchical structure of parental alleles to estimate the gene expression contributed by the parental strain (https://github.com/churchill-lab/emase). For each J:DO and gene, EMASE returns a vector with estimated gene expression values attributable to each of eight founder strains and the total gene expression, calculated as the sum of expression values across all eight founder strains. The proportion of gene expression contributed by each founder strain was calculated as the estimated gene expression values per founder strain divided by the total expression. EMASE was implemented through the Genotype by RNA-seq (GBRS) software package (https://gbrs.readthedocs.io/en/latest/; Choi et al, 2025).

### CellMAGE

Cell-type deconvolution for multi-parent population analysis of gene expression (CellMAGE) leverages known parental genetics to predict cell-type-specific gene expression in progeny from bulk RNA-seq data. The method requires two inputs: 1) snRNA-seq or scRNA-seq data from each parental strain, providing cell-type-specific expression profiles, and 2) bulk tissue allele-specific expression estimates for each progeny generated by EMASE from bulk RNA-seq (Fig. 1A).

**Figure 1.**
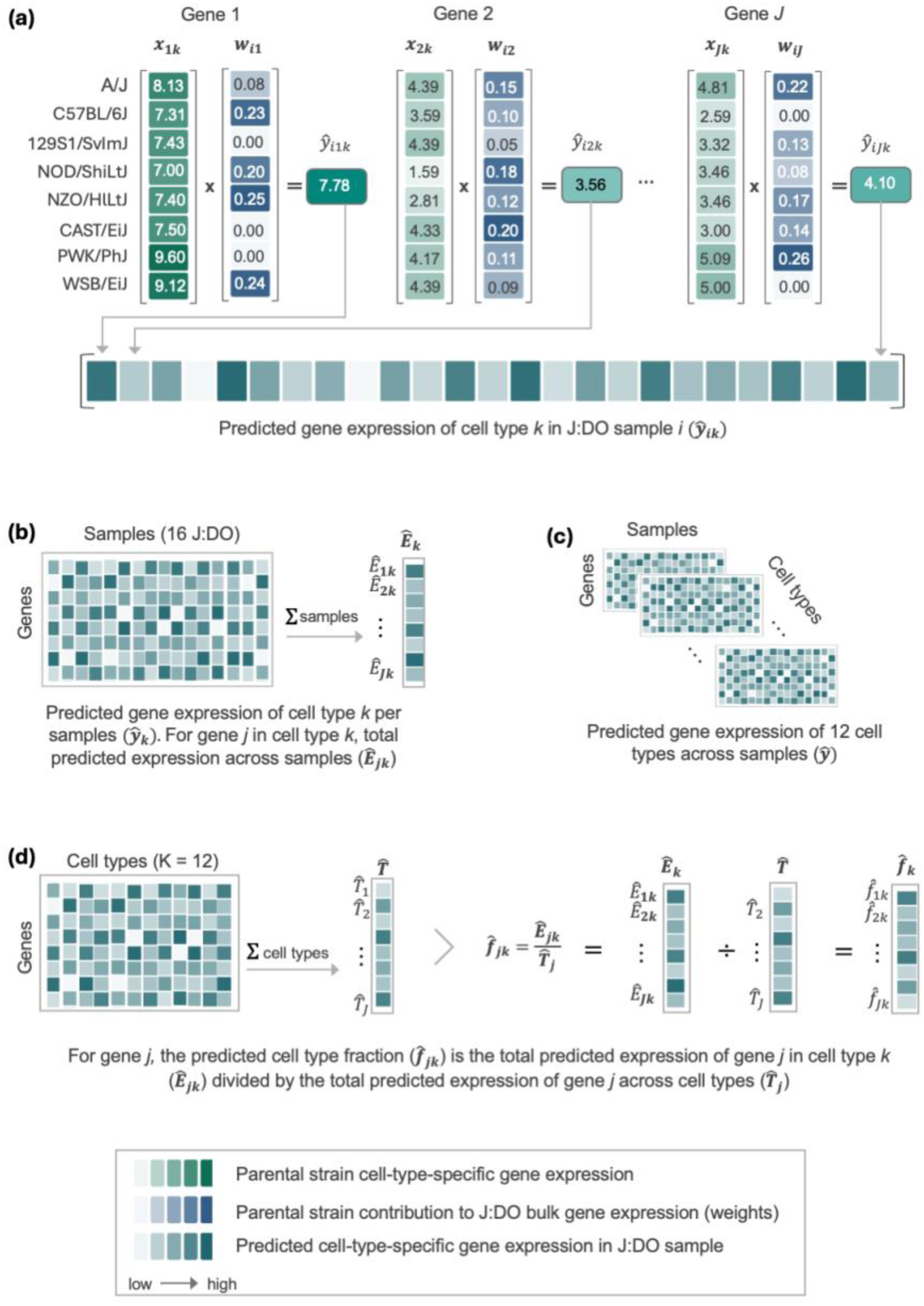
(**a**) CellMAGE predicts cell-type-specific expression (***ŷ_k_***; shown in teal) for gene *j* in a single J:DO sample (*i*) as the weighted sum of founder strain cell-type-specific gene expression (***x_jk_***; shown in green) and the EMASE-estimated relative contribution of the founder strain contribution to bulk expression for gene *j* in J:DO sample *i* (***w_ij_***; shown in blue). Cell-type-specific gene expression matrices (**b**) are predicted for all J:DO samples and across all cell types (**c**). Predicted cell type fractions (**d**) are calculated as the total predicted expression across marker genes for sample *i* in cell type *k* (***Ê_ik_***; panel **b**) divided by the total predicted expression for sample *i* across cell types (***T̂_i_***).

The proportional contribution of each parental strain to the gene expression in the progeny was calculated as EMASE estimated expression divided by the total expression. Consistent with the principle of parsimony, CellMAGE adopts the simplest model that fits the biological structure of the problem. Once each parent’s proportional contribution to the progeny bulk gene expression is known and parental cell-type expression profiles are measured, prediction reduces to a weighted sum. Weights reflect the proportional contribution of each parental strain to the gene expression, estimated from progeny bulk RNA-seq via EMASE (Fig. 1A). Sex-specific expression was estimated using cell-type expression profiles from parental strains of the same sex as the progeny.

Notably, CellMAGE does not require model training and therefore imposes no minimum sample size requirement. Predictions are generated per progeny sample using only that individual’s bulk RNA-seq data and parental cell-type expression profiles. No information is borrowed from other progeny. As a result, CellMAGE predicts cell-type-specific expression equally well for a single sample as for a large cohort.

The predicted cell-type expression (Fig. 1) is calculated as follows: Let *s* = 1, 2, … *S* parental strains, *i* = 1, 2, …, *N* progeny, *j* = 1, 2, … *J* genes, *k* = 1, 2, … *K* cell-types. The predicted expression *ŷ_ijk_* of gene *j* in cell-type *k* in progeny *i* (Fig. 1b & 1c) is estimated by the weighted sum of parental cell-type expression profiles *x_jks_*, where *w_ijs_* represents the proportional contribution of parental strain *s* to the estimated bulk gene expression in progeny *i*.

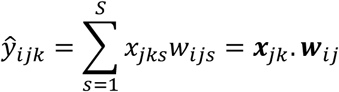

#### Predicted cell-type fraction

Predicted cell-type fractions were calculated per J:DO sample from predicted cell-type-specific gene expression. For each J:DO sample *i*, we first calculated the predicted abundance of cell-type *k* as the total cell-type-specific predicted expression across 116 marker genes, 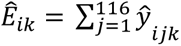 (Fig. 1b). The total predicted abundance across cell types for sample *i* was calculated as the sum of sample-level cell type abundances, 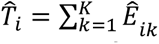. The predicted cell type fraction of cell type *k* in sample *i* was calculated as the proportion of predicted cell-type-specific abundance divided by total predicted abundance across cell types, 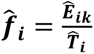, transforming the abundance to proportions so that the sum of predicted cell-type fractions within J:DO sample summed to one (Fig. 1d).

### CellMAGE Validation

CellMAGE predictions were validated against measured gene expression and cell-type fractions from individual snRNA-seq libraries per sample from the prefrontal cortex of 16 J:DO mice in a full factorial design (Table 1; Fig. 1b). CellMAGE performance was assessed by comparing predicted to measured cell type fractions within each J:DO sample and expression across 23,116 genes expressed in both founder cell-type-specific expression and J:DO bulk expression. For gene expression, cell-type-specific Spearman correlation and statistical equivalence was estimated per J:DO sample across genes (Fig. 2). Cell-type-specific equivalence of predicted and measured gene expression was tested with a linear mixed-effect model with random effects for gene and J:DO sample. Overall correlation and equivalence were assessed with mixed-effect models.

**Figure 2.**
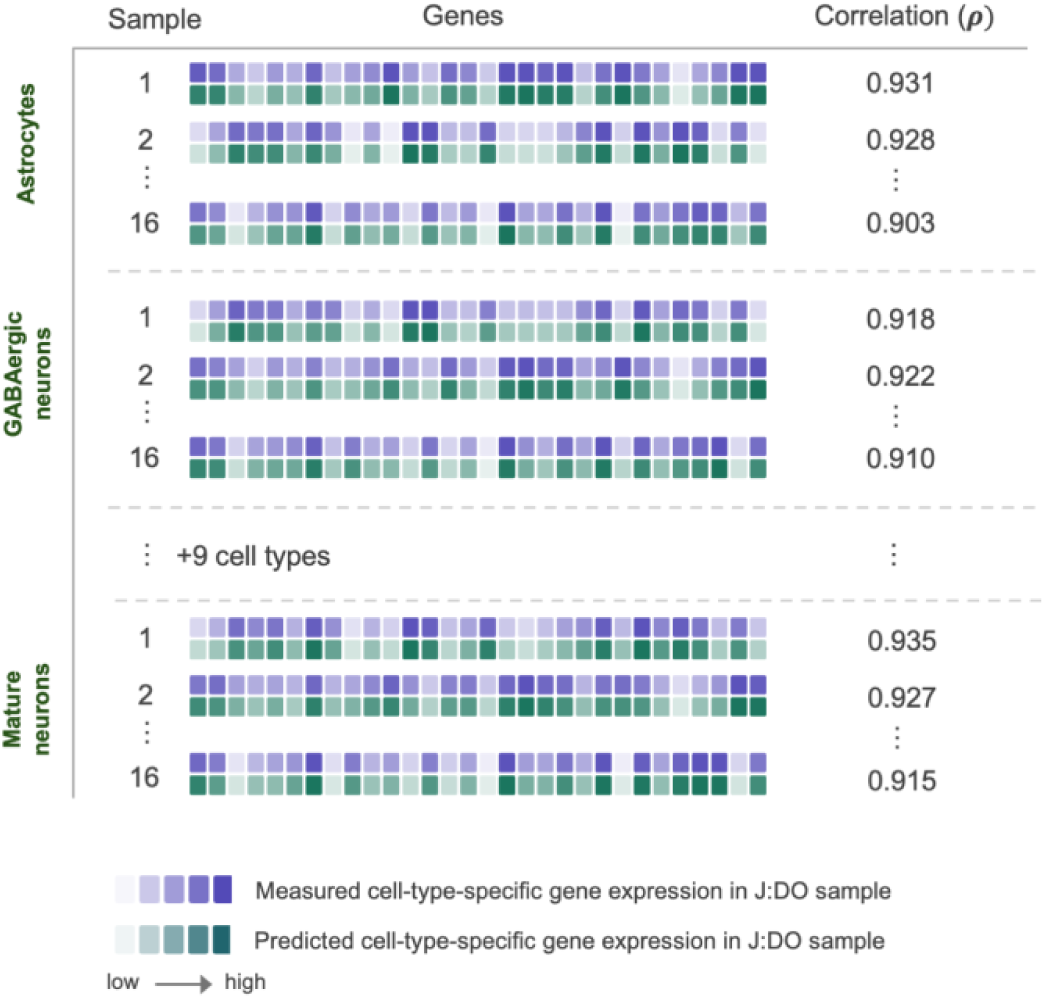
CellMAGE validation strategy. Predicted gene expression was validated against measured gene expression in 16 J:DO samples across 12 cell types. Spearman correlation (*ρ*) was calculated within sample and cell type across all genes expressed in the cell type, e.g., in astrocytes, each individual J:DO correlation was computed across >20,000 genes producing 16 correlation values all assessed across >20,000 genes. Equivalence within cell type was assessed with a mixed effect model with method (measured, predicted) as the fixed effect and random effects for sample and gene.

#### Spearman rank-order correlation: gene expression

Spearman correlation between predicted and measured gene expression was computed across cell types and J:DO samples and within cell type per J:DO sample. Specifically, for a cell-type and individual J:DO sample, Spearman’s *ρ* was calculated between predicted and measured gene expression for all genes expressed in the cell-type (Fig. 2). The overall Spearman’s *ρ* was estimated by pooling Spearman correlation values in a linear mixed-effect model, accounting for the correlation structure with random effects for J:DO sample and cell-type. Each of 192 (16 J:DO x 12 cell types) Spearman correlation values were Fisher-*z* transformed prior to modelling. The fixed effect intercept and profile likelihood 95% confidence interval from the mixed-effect model was back transformed via the hyperbolic tangent function (*tanh*) to obtain the pooled Spearman correlation and 95% confidence interval.

#### Spearman rank-order correlation: cell-type fraction

Agreement between measured and predicted cell-type fractions was assessed using Spearman’s ρ correlation, computed per sample across all cell types (12 cell types for both the primary CellMAGE analysis and the CellMAGE vs. CIBERSORTx fraction comparison). Each per-sample ρ was Fisher *z*-transformed, yielding a *z*-value with a known sampling variance based on the number of cell types included in each correlation. The resulting per-sample z-values were pooled via random-effects meta-analysis (rma function from metafor version 4.4-0, method = REML). The pooled z estimate and its 95% confidence interval were back-transformed via the hyperbolic tangent function (*tanh*) to obtain the overall Spearman correlation and its 95% confidence interval.

#### Equivalence test

Two one-sided tests (TOST) were used to assess statistical equivalence of predicted and measured values, evaluating whether the 90% confidence interval of the mean difference falls entirely within the predefined equivalence bounds, defined as the smallest effect size of interest (SESOI; Lakens, 2017). Equivalence bounds were set at ±0.05 for both gene expression and cell-type fractions. To correct for multiple comparisons across cell types, Benjamini-Hochberg (BH) correction was applied to equivalence test *p*-values for both gene expression and cell-type fraction analyses.

Gene expression was *z*-score normalized prior to analysis. Equivalence of predicted and measured gene expression was assessed within each cell type using a linear mixed-effects model with method (predicted vs. measured) as the fixed effect and random effects for J:DO sample and gene. To obtain a pooled equivalence estimate across cell types, cell-type-specific equivalence estimates and standard errors were combined in a random-effects meta-analysis using restricted maximum likelihood (REML; rma function, metafor package v4.4-0; Viechtbauer, 2010), which accounts for both within-cell-type sampling error and between-cell-type heterogeneity (τ²). A 90% confidence interval was computed as estimate ± t(0.95, df = k−1) × SE, where k is the number of cell types, and evaluated against the ±0.05 equivalence bounds using TOST. Between-cell-type heterogeneity was assessed using the Q-test, I², and τ² to confirm that the random-effects approach was appropriate. Equivalence of predicted and measured cell-type fractions was assessed by directly computing paired differences between predicted and measured proportions per sample, which were then pooled across cell types using the same random-effects meta-analytic approach described above.

#### Sensitivity Analyses

CellMAGE predictions are based on two inputs: parental cell-type-specific expression profiles and progeny bulk gene expression, both of which involve preprocessing choices that could in principle affect prediction accuracy. Normalization strategy and pseudobulk construction method have both been shown to substantially affect the accuracy of reference-based deconvolution methods (Xu et al., 2025). To assess the robustness of CellMAGE predictions under alternative preprocessing approaches, we conducted sensitivity analyses examining whether performance improved when founder snRNA-seq data were normalized jointly with 8 saline-treated J:DO samples, and whether performance was sensitive to pseudobulk construction method.

Founder expression profiles were normalized and preprocessed independently from the J:DO samples. We evaluated whether joint normalization, incorporating 8 saline-treated J:DO samples (4 per sex) into the founder preprocessing pipeline, improved prediction performance. All other pipeline inputs and preprocessing steps were held constant.

Additionally, founder cell-type expression profiles were constructed as the sum of counts across cells per sample per cell type using Seurat’s AggregateExpression() function. To assess sensitivity to pseudobulk construction method, we evaluated whether prediction performance changed when founder cell type expression profiles were constructed as the mean of counts across cells per sample per cell type, using Seurat’s AverageExpression() function. All other inputs were held constant. Predictions were validated against J:DO measured expression, also constructed with AverageExpression().

As described above (Methods; CellMAGE validation), CellMAGE performance was assessed with the equivalence test and Spearman correlation.

### Software availability

CellMAGE is freely available as an R Shiny application for interactive use without programming expertise and as an R package for integration into large-scale computational pipelines, with full documentation and example datasets at https://thejacksonlaboratory.shinyapps.io/CellMAGE/ and upon acceptance, at https://github.com/TheJacksonLaboratory/chesler-lab/tree/main/CellMAGE. To demonstrate applicability beyond mammalian multiparental populations, application documentation includes a worked example applying CellMAGE to a multiparental maize (*Zea mays*) population (Miculan et al., 2021; Supplementary Methods), with input data and annotated code provided to facilitate adaptation to other species and population designs (see Web Resources).

### Comparative analysis

CellMAGE was compared to CIBERSORTx (Newman et al., 2019), a widely used deconvolution method and benchmark in the field. Both methods used founder snRNA-seq data as the cell-type reference, CellMAGE as cell-type-specific expression profiles and CIBERSORTx as the signature matrix. CIBERSORTx predicted cell-type fractions and gene expression for 112 J:DO bulk RNA-seq samples, the 16 validation samples plus an additional 96 J:DO samples balanced for sex and treatment (Table 1). The expanded cohort was required to meet CIBERSORTx’s minimum sample size requirement for cell-type-specific gene expression estimation, which requires 3–4 times more samples than cell types. Both methods were evaluated against measured cell-type transcriptomes from the 16 J:DO validation samples.

#### CIBERSORTx predicted cell type fractions

CIBERSORTx cell-type fractions were estimated for all 12 cell types using ‘Cell Fractions’ mode with 1,000 permutations and quantile normalization disabled. Equivalence testing and Spearman correlation were calculated as described above, with one exception: Fisher-*z* transformation is undefined at ρ = ±1, which can occur with only 12 cell types per sample. In such cases, ρ was capped at ±0.999 prior to transformation and mixed-effects meta-analysis.

CIBERSORTx requires a single-cell reference as input when data were generated using a UMI-based platform such as 10x Genomics. This reference is used to construct a signature matrix for estimating cell-type fractions, total gene expression profiles, and cell-type-specific gene expression. Because single-cell UMI count data and bulk RNA-seq differ systematically in expression distributions and technical noise, a signature matrix built directly from single-cell data may not translate directly to bulk RNA-seq expression space. For this reason, instead of using pseudobulk as input, we used single cell reference profiles with CIBERSORTx’s S-mode batch correction, which is recommended for signature matrices derived from droplet-based, UMI-based scRNA-seq platforms such as 10x Chromium.

The single-cell reference was constructed by randomly sampling 50% of cells per cell type across all 12 cell types (sampling = 0.5), yielding 132,751 cells total. Genes were included if there was a minimum of 75% of cells within a cell type to show evidence of expression (fraction = 0.75; Supplementary Data S1). Cell-type labels were inferred directly from the reference without explicit phenotype class assignment. Reference profiles were constructed using 5 pseudo-replicates per cell type (replicates = 5), with signature matrix gene selection requiring a minimum of 300 and maximum of 500 genes per cell type (G_min = 300, G_max = 500) and a differential expression *q*-value ≤ 0.01. Matrix conditioning was controlled with a maximum condition number of 999 (k.max = 999). The resulting unadjusted signature matrix (n = 2,732 genes; Supplementary Data S1) was uploaded alongside the adjusted reference sample to the CIBERSORTx web interface (https://cibersortx.stanford.edu, accessed August 20, 2026). The mixture file comprised 112 J:DO bulk RNA-seq samples (Table 1). S-mode batch correction was applied to address platform differences between the single-cell UMI reference and bulk RNA-seq mixture data, as recommended for droplet-based UMI platforms such as 10x Chromium.

For the comparative analysis, CellMAGE and CIBERSORTx cell-type fraction predictions were based on 2,732 genes in the CIBERSORTx signature matrix. Predicted cell-type fractions were compared to the 16 J:DO validation samples using the same Spearman correlation and TOST equivalence testing framework described above.

#### CIBERSORTx predicted gene expression (High Resolution)

Cell-type fraction results (Supplementary Data S1) were used as input (--cibresults) to CIBERSORTx’s High Resolution (HiRes) mode, run via Singularity container (HiRes, latest version downloaded January 10, 2025). Using the same unadjusted signature matrix and adjusted reference sample described above (--sigmatrix, --refsample; built from founder mouse reference data), HiRes mode was run across all 112 J:DO bulk (mixture) samples for all 12 cell types in the signature matrix (24,893 genes), with an adaptive window size of 48 (default value, calculated as number of cell types x 4 = 48). S-mode batch correction was applied (--rmbatchSmode TRUE) to account for technical differences between the single-cell-derived reference and bulk mixture data. HiRes does not itself take a permutation parameter; instead, prior CIBERSORTx Fractions results (--cibresults, run with 1,000 permutations) were supplied to inform per-sample cell-type composition during HiRes expression imputation.

To ensure a fair comparison, CellMAGE predicted cell-type specific expression was restricted to the same genes retained after CIBERSORTx filtering. Both methods were then evaluated against the 16 J:DO validation samples using Spearman correlation and TOST equivalence testing, as described above.

Except where noted, all analyses were performed in the R environment for statistical computing, version 4.3.1 (R Core Team, 2023).

## Results

CellMAGE predicted cell-type-specific gene expression for 16 J:DO PFC samples across 12 cell types: astrocytes, dopaminergic neurons, endothelial cells, GABAergic neurons, glutamatergic neurons, mature neurons, microglial cells, neural progenitor cells, neuroepithelial cells, oligodendrocyte precursor cells, oligodendrocytes, and tanycytes. Gene coverage ranged from 20,044 to 22,876 genes per cell type, total 23,116 genes (Supplementary Data S2). Predicted and measured gene expression were highly correlated, with a pooled Spearman correlation of 0.923 (95% CI: 0.910, 0.934; Fig. 3a), and statistically equivalent (±0.05) in all 12 cell types after BH correction (Fig. 3c). Predicted cell-type fractions were also accurate, with a pooled Spearman correlation of 0.826 (95% CI: 0.765, 0.872; Fig. 3b) and statistical equivalence achieved in 7 of 12 cell types after BH correction (Fig. 3d; Supplementary Data S2).

**Fig 3.**
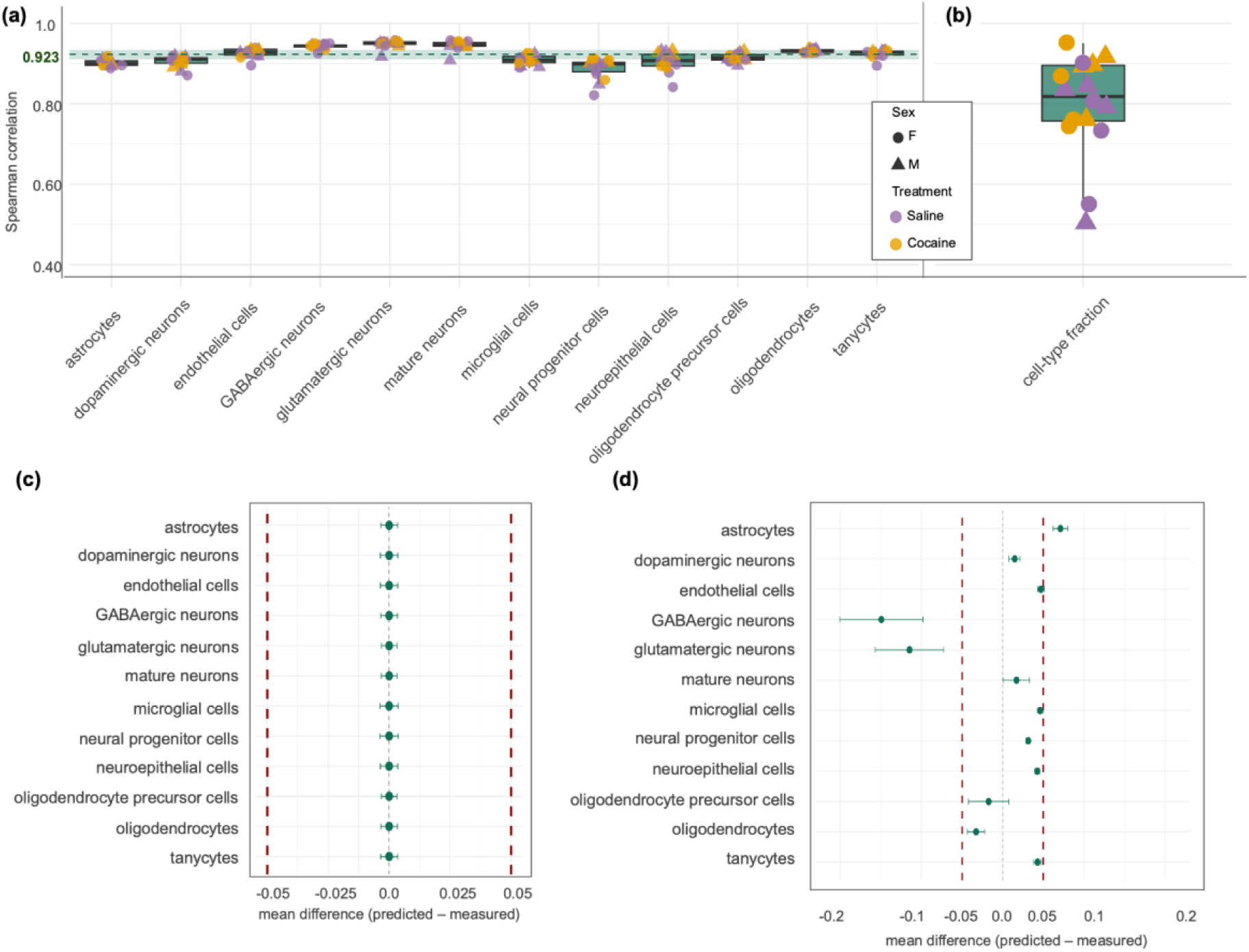
Validation of CellMAGE-predicted vs measured gene expression and cell-type fractions. **(a)** Spearman correlation between CellMAGE predicted and measured gene expression per sample within cell type, and (**b)** predicted vs measured cell-type fractions across J:DO samples. Each point represents one J:DO sample (n = 16 per cell type), colored by treatment (saline = purple, cocaine = orange) and shaped by sex (female = circle, male = triangle). Boxplots show the median and interquartile range across samples. Dashed line indicates the pooled Spearman correlation (ρ = 0.923), shaded area indicates pooled 95% CI (0.910, 0.934). **(c)** Equivalence test results for CellMAGE-predicted vs. measured expression by cell type and **(d)**

Sensitivity analyses confirmed robustness to input preprocessing. In the primary analysis, founder data was normalized independently of saline-treated J:DO samples and pseudobulk was constructed by aggregating expression. Normalizing the founder data with saline-treated J:DO samples and constructing pseudobulk with average expression resulted in pooled Spearman correlations of 0.931 (95% CI: 0.920, 0.940) and 0.930 (95% CI: 0.918, 0.940), respectively. Pooled Spearman correlation is similar to the primary analysis – 0.923 (95% CI: 0.910, 0.934) – with overlapping confidence intervals. Equivalence tests across all 12 cell types were comparable to the primary analysis under both alternative approaches (Supplementary Data S3). CellMAGE also outperformed CIBERSORTx in cell-type fraction concordance, achieving a substantially higher pooled Spearman correlation (0.808 [95% CI: 0.738, 0.861] vs. 0.607 [95% CI: 0.494, 0.701]) and substantially higher gene expression concordance across all three comparable cell types (median ρ ranging from 0.908 to 0.974 vs. 0.353 to 0.641). Notably, CIBERSORTx resolved gene expression in only 3 of 12 cell types and at most 3,084 of 23,116 (13.3%) genes per cell type, compared to CellMAGE which predicted expression for all 20,000+ genes across all 12 cell types (Supplementary Data S4).

### Validation of CellMAGE predicted gene expression

Predicted and measured gene expression were compared across 20,044–22,876 genes per cell type. In all 12 cell types, predicted and measured expression were highly correlated, with median Spearman correlation ranging from 0.899 to 0.951 (Fig. 3a; Supplementary Data S2). Concordance was consistent across all 16 samples (IQR < 0.05 for all cell types), indicating that variation in Spearman correlation was small across treatment and sex groups. Pooled across cell types, the Spearman correlation was 0.923 (95% CI: 0.910, 0.934).

Predicted gene expression was statistically equivalent to measured expression (±0.05) in all 12 cell types after BH correction (Fig. 3c). Mean differences were negligible across all cell types (range: −3.78 × 10⁻¹⁷ [microglial cells, 90% CI: −3.83 × 10⁻³ to 3.83 × 10⁻³] to 2.26 × 10⁻¹⁷ [oligodendrocyte precursor cells, 90% CI: −3.48 × 10⁻³ to 3.48 × 10⁻³], median: −7.67 × 10⁻¹⁸). Pooled across cell types, predicted and measured gene expression were statistically equivalent (±0.05), with a mean difference of approximately zero (90% CI: −0.001, 0.001; TOST *p* < 0.001; Supplementary Data S2).

### Validation of CellMAGE predicted cell-type fractions

CellMAGE achieved high concordance with measured cell-type fractions across all 12 cell types (pooled Spearman ρ = 0.826 [95% CI: 0.765, 0.872]). For each sample, a single Spearman ρ was calculated across its 12 cell types, yielding one correlation per sample; these 16 per-sample correlations were also high (median [IQR] = 0.818 [0.757, 0.895]; Fig. 3b; Supplementary Data S2), and were pooled via random-effects meta-analysis to obtain the overall estimate above. Between-sample heterogeneity was negligible, indicating that the pooled estimate was not meaningfully driven by differences across samples (τ² = 0.005 [95% PL CI: 0.000, 0.150]; I² = 4.1% [95% CI: 0.0, 57.4%]).

CellMAGE predicted cell fractions were statistically equivalent to measured cell fractions (±0.05 across 116 marker genes; see Methods) in 7 of 12 cell types after BH correction, including dopaminergic neurons, mature neurons, neural progenitor cells, neuroepithelial cells, oligodendrocyte precursor cells, oligodendrocytes, and tanycytes. Astrocytes, endothelial cells, GABAergic neurons, and glutamatergic neurons did not achieve statistical equivalence. Microglial cells narrowly missed the equivalence threshold (BH-adjusted *p* = 0.054; Fig. 3d).

### Sensitivity Analyses

CellMAGE predictions depend on two inputs: parental cell-type-specific expression profiles and progeny bulk gene expression, both of which involve preprocessing choices that could in principle affect prediction accuracy. Normalization strategy and pseudobulk construction method have both been shown to substantially affect the accuracy of reference-based deconvolution methods (Xu et al., 2025). To assess the robustness of CellMAGE predictions to these choices, we conducted two sensitivity analyses: (1) founder expression profiles normalized jointly with saline-treated J:DO samples rather than independently, and (2) pseudobulk expression constructed using AverageExpression() instead of AggregateExpression() for both founder profiles and J:DO validation data.

In both analyses, predicted and measured gene expression were highly correlated, with pooled Spearman correlations of 0.931 (95% CI: 0.920, 0.940) for the joint normalization analysis and 0.930 (95% CI: 0.918, 0.941) for the mean expression analysis, both marginally higher than the primary analysis (ρ = 0.923, 95% CI: 0.910, 0.934). Between-cell-type heterogeneity was negligible in both analyses (I² = 0%, τ² = 0), indicating that pooled estimates were not meaningfully driven by variability across cell types. Predicted gene expression remained statistically equivalent to measured expression (±0.05) in all 12 cell types after BH correction under both alternative approaches, with negligible mean differences across cell types (all < 0.001). Per-cell-type equivalence test and correlation results are reported in Supplementary Data S3.

Results were consistent across both sensitivity analyses, supporting the robustness of CellMAGE predictions to input preprocessing choices.

### Comparative Analysis

#### CIBERSORTx gene expression resolution

CIBERSORTx returned predicted expression values for 7,342 of 23,177 (31.7%) genes in the validation set; the remainder were assigned a value of 1, denoting insufficient evidence of expression (CIBERSORTx HiRes README, Newman Lab, https://cibersortx.stanford.edu, accessed August 20, 2026). Though not included in the final comparison, CIBERSORTx returned gene expression predictions for 12 cell types. To ensure a meaningful comparison against measured data, cell types were retained for downstream analysis only if CIBERSORTx returned usable predicted expression for at least 500 genes across all 16 validation samples. After filtering, three cell types met this criterion: glutamatergic neurons, microglial cells, and oligodendrocytes. All predictions returned by CIBERSORTx are available in Supplementary Data S1.

#### Comparison of CellMAGE and CIBERSORTx predicted gene expression

Despite statistically equivalent mean differences, the two methods differed meaningfully in concordance between predicted and measured expression. CellMAGE achieved high concordance across all three cell types (pooled Spearman ρ = 0.944 [95% CI: 0.861, 0.978]), with median Spearman correlations ranging from 0.908 to 0.974 compared to CIBERSORTx (pooled Spearman ρ = 0.552 [95% CI: 0.325, 0.719]), with median Spearman correlations that ranged from 0.353 to 0.641; a median difference greater than 0.582 (Fig. 4a). Notably, CIBERSORTx predicted expression in microglial cells was particularly poor, with a median Spearman correlation of 0.353. Correlation estimates were consistent across the 16 J:DO samples for both methods, with interquartile ranges below 0.05 in all cases (Supplementary Data S4), indicating that the performance gap was systematic rather than driven by a subset of samples.

**Fig 4.**
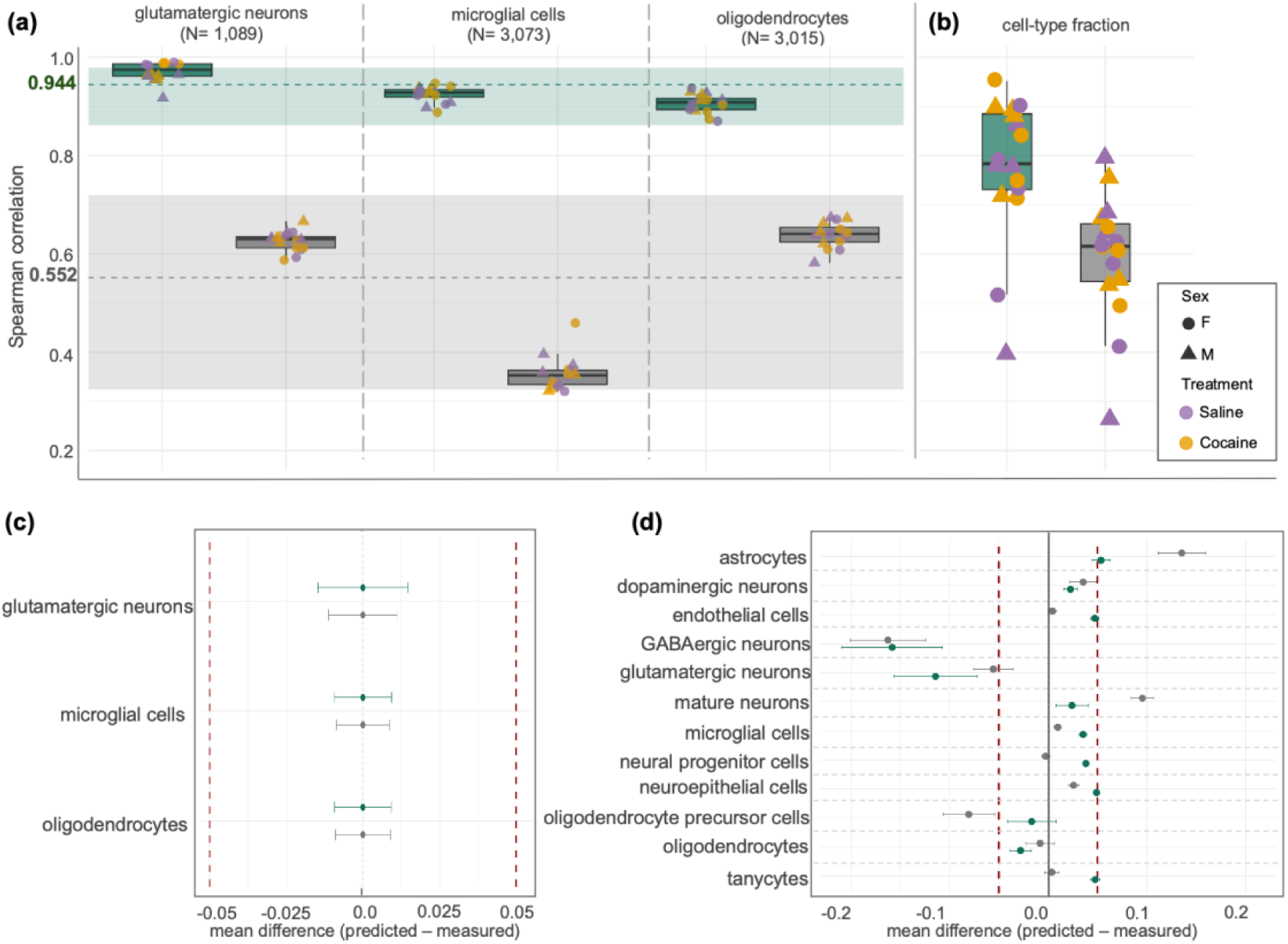
Comparative analysis validation of CellMAGE and CIBERSORTx. (**a, b)** Spearman correlation between predicted and measured expression **(a)** for each cell type with the number of genes listed below and cell type fractions **(b)**. Each point represents one J:DO sample in the validation set (n = 16), colored by treatment (saline = purple, cocaine = orange) and shaped by sex (female = circle, male = triangle). Boxplots show median and interquartile range across samples. Dashed line indicates the pooled Spearman correlation with 95% CI shaded area. Forest plots of the mean difference (predicted − measured) for CellMAGE (teal) and CIBERSORTx (gray) across cell-type expression **(c)** and cell-type fractions **(d)** using the genes in the CIBERSORTx signature matrix. Points represent mean differences; error bars indicate 90% confidence intervals. Dotted red lines denote equivalence bounds (±0.05); cell types with confidence intervals falling entirely within these bounds are considered statistically equivalent to measured values.

Both methods achieved statistical equivalence (±0.05) in predicted gene expression in all three cell types after BH correction (Fig. 4c; Supplementary Data S4), with negligible mean differences across cell types for both methods (CellMAGE: range −1.11 × 10⁻¹^8^ to 1.07 × 10⁻¹⁷, median 4.27 × 10⁻¹^8^; CIBERSORTx: range −8.32 × 10⁻¹⁷ to 1.97 × 10⁻¹⁵, median −6.89 × 10⁻¹⁷). Pooled across the three cell types, overall statistical equivalence was confirmed for CIBERSORTx (90% CI: −0.0097, 0.0097; TOST *p* = 0.0022) and approached for CellMAGE (90% CI: −0.011, 0.011; TOST p = 0.0026), with no evidence of heterogeneity across cell types for either method (τ = 0, I² = 0%, Q-test p = 1).

#### CIBERSORTx cell-type fraction resolution

CIBERSORTx returned deconvolution results for all 16 J:DO validation samples (Supplementary Data S1) across 2,732 genes. CIBERSORTx internal quality metrics indicated a moderate fit with a mean correlation of 0.516 (SD = 0.070; range: 0.399–0.635) and RMSE of 0.856 (SD = 0.041; range: 0.783–0.924).

All 12 cell types were represented across the sample set, however, CIBERSORTx returned zeroes for some samples within specific cell types. Oligodendrocyte precursor cells were most affected, with predicted cell fraction = 0 in 7 of 16 samples (44%), suggesting this cell type may be at or near the detection limit of the CIBERSORTx signature matrix.

#### Comparison of CellMAGE and CIBERSORTx predicted cell type fractions

CellMAGE achieved substantially greater concordance with measured cell-type fractions than CIBERSORTx across all 12 cell types (pooled Spearman ρ = 0.808 [95% CI: 0.738, 0.861] vs. 0.608 [95% CI: 0.494, 0.701]), a difference of 0.200 correlation and non-overlapping confidence intervals (Fig. 4b). Per-sample correlations were also higher for CellMAGE (median [IQR] = 0.783 [0.731, 0.885]) than CIBERSORTx (median [IQR] = 0.615 [0.544, 0.661]; Fig. 4b; Supplementary Data S4). Between-sample heterogeneity was negligible for both methods, indicating that pooled estimates were not meaningfully driven by differences across samples (CellMAGE: τ² = 0.018 [95% PL CI: 0.000, 0.179], I² = 13.7% [95% CI: 0.0, 61.7%]; CIBERSORTx: τ² = 0 [95% PL CI: 0–0.048], I² = 0% [95% CI: 0.0, 30.2%]).

CellMAGE and CIBERSORTx predicted and measured cell-type fractions were statistically equivalent (±0.05) in 6 of 12 cell types after BH correction, with both methods resolving microglial cells, neural progenitor cells, and oligodendrocytes (Fig. 4d). CIBERSORTx uniquely resolved endothelial cells, neuroepithelial cells, and tanycytes, while CellMAGE uniquely resolved dopaminergic neurons, mature neurons, and oligodendrocyte precursor cells. Predicted fractions for astrocytes, glutamatergic neurons, and GABAergic neurons did not achieve statistical equivalence to measured fractions under either method, however, mean differences spanned a narrower range for CellMAGE (−0.158 to 0.053; 90% CI bounds: −0.209 to 0.062) than for CIBERSORTx (−0.163 to 0.135; 90% CI bounds: −0.201 to 0.159), suggesting more consistent cell-type fraction estimates for CellMAGE.

## Discussion

CellMAGE successfully exploits the known genetic architecture of multiparental populations to reconstruct cell-type-specific transcriptomes in progeny from bulk RNA-seq data. The central premise is that each progeny’s transcriptome reflects a quantifiable mixture of parental transcriptomes, a structure that provides a principled biological basis for deconvolution without model fitting. Rather than inferring cellular composition from statistical patterns in the data, CellMAGE derives predictions directly from parental genetics, weighting parental cell-type profiles by each progeny’s EMASE-estimated allele-specific expression.

CellMAGE achieved excellent prediction accuracy across all 12 prefrontal cortex cell types, with predicted gene expression statistically equivalent to measured expression (±0.05) and a pooled Spearman correlation of 0.923 (95% CI: 0.910, 0.934). Performance was demonstrated across a full factorial sex-by-treatment design, establishing that CellMAGE recovers biologically meaningful variation driven by sex and environment, as evidenced by negligible between-sample heterogeneity across groups. Normalization strategy and pseudobulk construction method are known to substantially affect the accuracy of reference-based deconvolution methods (Xu et al., 2025), but because CellMAGE computes predictions as a weighted sum of parental profiles rather than fitting parameters to the reference, these preprocessing choices were not expected to substantially alter performance, an expectation confirmed by both sensitivity analyses.

CIBERSORTx is a well-validated deconvolution method that performs well in conventional population studies but was designed for settings where the genetic relationship between reference and target is unknown. In multiparental populations this inference is unnecessary, and CIBERSORTx’s model-based approach imposed a minimum sample size requirement necessitating 112 bulk RNA-seq samples compared to 16 for CellMAGE. Despite both methods passing equivalence testing for mean differences, CellMAGE achieved markedly higher concordance with measured expression (per-cell-type median ρ: 0.908–0.974 vs. 0.353–0.641) and resolved all 23,116 genes across all 12 cell types, compared to at most 3,084 genes (13.3%) in 3 of 12 cell types for CIBERSORTx.

Several limitations should be noted. Prediction accuracy depends on the alignment between parental reference profiles and the tissue being deconvolved; non-tissue-specific profiles would be expected to degrade performance. Non-additive genetic effects could in principle reduce accuracy, though negligible variation across treatment and sex groups suggests these did not meaningfully impact results. The failure to achieve cell-type fraction equivalence for GABAergic and glutamatergic neurons likely reflects the transcriptional similarity of these populations and growing evidence that many neurons co-express markers for both neurotransmitters (Brunet Avalos & Sprecher, 2021), a limitation of cell-type annotation rather than of the deconvolution method, as evidenced by CIBERSORTx’s equivalent failure for these same cell types. Finally, CellMAGE was benchmarked against CIBERSORTx alone; comparison to additional deconvolution tools would provide a more complete performance picture.

Beyond the Diversity Outbred mouse population studied here, CellMAGE is directly applicable to any multiparental population with matched parental single-cell data, including Collaborative Cross mice, heterogeneous stock rats, *Drosophila*, *C. elegans*, *Arabidopsis*, and diploid crop MAGIC populations. As demonstrated in a companion maize example (Miculan et al., 2021; Supplementary Methods), the method requires no modification when applied to non-mammalian species. Extension to polyploid crop species such as wheat will require adaptation of the allele-specific expression framework underlying EMASE to accommodate multi-copy subgenomic structures, and application to multi-breed livestock composite populations will become feasible as parental breed-specific single-cell atlases expand, though breed-specific variant catalogs such as the 1000 Bull Genomes Project (Hayes & Daetwyler, 2019) already provide the genomic infrastructure required for EMASE alignment. Because CellMAGE requires no model fitting, it imposes no minimum sample size. Predictions are generated per individual from the individual’s bulk RNA-seq data and the parental reference profiles, with no information borrowed across samples. This makes CellMAGE particularly valuable for pilot studies, rare disease models, single-animal experiments, and retrospective application to existing bulk RNA-seq datasets from multiparental populations originally collected without single-cell profiling in mind. The parental single-cell reference is a one-time investment *per tissue*, not *per study* — once founder profiles are established, every subsequent bulk RNA-seq study in that tissue gains cell-type resolution at bulk sequencing cost. As reference genomes and single-cell atlases continue to expand across organisms, the scope of CellMAGE’s applicability will grow accordingly.

CellMAGE is freely available as an R Shiny application for interactive use and an R package for large-scale pipelines, with full documentation and example datasets (see Web Resources).

### Web resources

CellMAGE is freely available as an R Shiny application and R package at https://thejacksonlaboratory.shinyapps.io/CellMAGE/ and upon acceptance, at https://github.com/TheJacksonLaboratory/chesler-lab/tree/main/CellMAGE/. The Shiny application supports interactive deconvolution and result exploration without programming expertise, accepting formatted input data and returning a downloadable table of predicted cell-type-specific gene expression alongside a gene-level visualization module. The R package is recommended for computational pipelines or studies with input data exceeding 3 GB, typical of larger cohorts. Documentation includes user guides and example datasets for mouse and maize populations (Supplementary Data S5).

**Fig 5.**
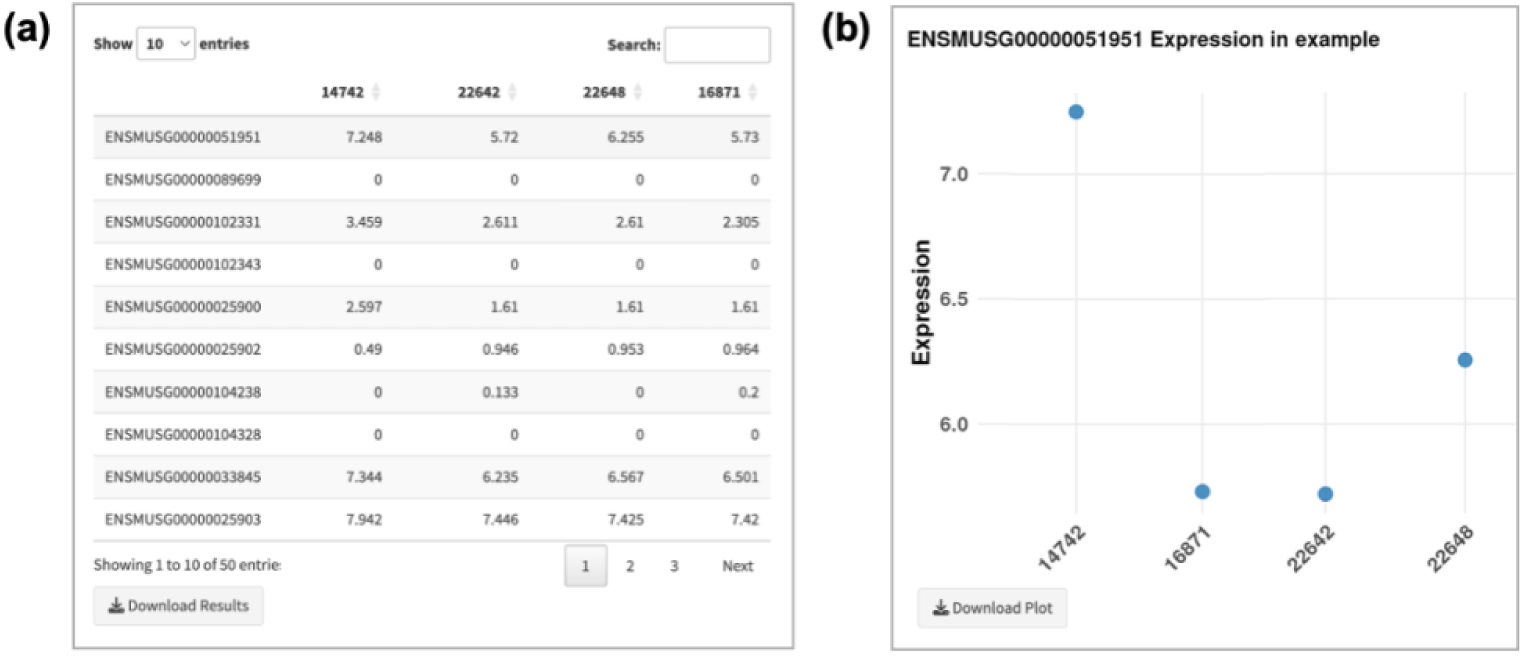
CellMAGE R Shiny Application. **(a)** The CellMAGE R Shiny app returns a downloadable table of predicted cell-type-specific gene expression results. **(b)** Users also have the option to use the gene visualization module, where users can select a gene of their choice and specific samples to explore predicted expression at the gene level.

## Data availability statement

Raw sequencing data (bulk RNA-seq and snRNA-seq) have been deposited in the NCBI Sequence Read Archive (SRA; accessions PRJNA1517767, PRJNA1516495, PRJNA1518853). Processed expression data have been deposited in Gene Expression Omnibus (GSE346204).

Supplemental Data is available at Figshare. Each folder contains a README with detailed information for each file included. Folder S1 contains all material relevant to CIBERSORTx, including the single cell reference, refsample, and signature matrix constructed to run the pipeline, predicted gene expression, and predicted cell-type fractions. Folder S2 contains all relevant material for CellMAGE validation analyses, including all supporting files for calculating predicted cell-type fractions, single cell metadata, gene counts per cell type, predicted cell-type fractions, predicted gene expression, and validation testing results (Spearman rank correlation and equivalence test). Folder S3 includes all material relevant for CellMAGE sensitivity analyses, including gene counts per cell type, predicted gene expression, and validation testing results (for each sensitivity analysis-joint normalization and average expression). Folder S4 contains all files for comparative analyses between CellMAGE and CIBERSORTx, including metadata for additional samples needed to run CIBERSORTx, filtered expression data based on usable gene expression from CIBERSORTx HiRes output, and all validation test results per method (Spearman rank correlation and equivalence test). Folder S5 contains Zea Mays example data, including inputs required for running CellMAGE.

The custom scripts and source code used in this study will be made publicly available on GitHub at https://github.com/TheJacksonLaboratory/chesler-lab/tee/main/CellMAGE/CellMAGE upon peer-review acceptance and publication. Currently, all code is available in Figshare for ease of review. Code-singlecell contains all code needed to generate the Seurat objects of the single cell data used in the analyses. Code-preprocessing contains all code to construct measured cell-type fractions using the measured Diversity Outbred (J.DO) mouse expression, calculate the predicted cell-type fractions based on the predicted gene expression from CellMAGE, and process the measured data with the predicted data so the appropriate sets of genes were used for each cell type in the primary analysis, as well as the sensitivity analyses. Code-spearman includes all code used to calculate Spearman rank correlation for expression as well as cell-type fraction for the CellMAGE primary analysis (S2), sensitivity analyses (S3), and comparative analyses (S4). Code-equivalence_test includes all code used to set up and run TOST equivalence tests for all expression and cell-type fraction analyses, for the CellMAGE primary analysis (S2), sensitivity analyses (S3), and comparative analyses (S4). Code-maize_example includes all code used to process EMASE files for the maize bulk samples, as well as code for building the maize-based inputs for CellMAGE in the required format. Code-CellMAGE includes all code in the build of the CellMAGE R package. An R Shiny app of CellMAGE is also available at https://thejacksonlaboratory.shinyapps.io/CellMAGE/.

## Acknowledgements

We gratefully acknowledge the contributions of Sandy Daigle, Elise Curtois, Mary Barter, and the JAX Single Cell Biology and Genome Technologies Service at The Jackson Laboratory. We thank Michael Lloyd of JAX Data Science for his assistance with the *Zea Mays* example and Ruohan Wang and Andrew Gentles of Stanford University for their assistance in running CIBERSORTx.

## Study funding

This work was supported by NIDA R01 DA037927 (Chesler) and P50 DA039842 (CSNA), with support from The Jackson Laboratory (JAX) Scientific Services Innovation Fund (Skelly) and NCI CCSG P30CA034196.

## Conflict of interest

No conflicts of interest to report.

## Author contributions

R.L.B. and E.J.C. conceptualized the project. A.A.A collected tissue with oversight from L.G. R.L.B. developed CellMAGE, the cell deconvolution statistical approach and algorithm. A.K. and R.L.B. validated CellMAGE. A.K. ran CIBERSORTx and compared CellMAGE and CIBERSORTx predictions. A.A.A and D.S. analyzed raw snRNA-seq single cell data. H.H. and V.M.P. analyzed raw bulk RNA-seq data and in collaboration with A.A.A., curated raw and processed data into the Short Read Archive. A.K. developed the CellMAGE R Shiny app and R package with oversight from R.L.B. A.A.A., A.K., R.L.B., and E.J.C. wrote the paper with input from all authors.

